# A 5′-end based Molecular Marker Enables Identification and Deployment of a Functional *Rpi-vnt1.1* Allele for Durable Late Blight Resistance in Potato

**DOI:** 10.1101/2025.11.06.687089

**Authors:** Shuo Wang, Luyao Wang, Simeng Feng, Chenxi Hu, Xi Meng, Shiwei Chang, Yan Feng, Liang Kong, Xiaoqiang Zhao

## Abstract

Potato (*Solanum tuberosum*) is a globally important food crop, yet its production is severely threatened by late blight, caused by *Phytophthora infestans*. Deploying resistance (*R*) genes in breeding programs offers a sustainable approach for disease management. *Rpi-vnt1.1* is a broad-spectrum *R* gene cloned from wild *Solanum* species *Solanum venturii*. Here, we developed a PCR marker targeting the 5′-end extension of *Rpi-vnt1.1*, which enables efficient and specific detection of the resistant alleles. Screening of advanced breeding lines from the International Potato Center (CIP) identified line C1848, which carries a functional *Rpi-vnt1.1* allele. Inoculation assays confirmed C1848 confers strong resistance to *P. infestans*. Subsequent functional assays, including RNAi-mediated silencing in C1848 and heterologous expression in *Nicotiana benthamiana*, demonstrated that this resistance is specifically mediated by the *Rpi-vnt1.1* allele. This specific marker provides a rapid, low-cost tool for marker-assisted breeding, facilitating the deployment of potato cultivars carrying *Rpi-vnt1.1*.

---

Potato (*Solanum tuberosum* L.) is one of the world’s most important food crops, ranking third globally in terms of production after rice and wheat (Haverkort et al., 2009). It is a staple food in many regions, providing a significant source of nutrition due to its high carbohydrate content (Beals, 2019). Potato cultivation spans across Asia, Europe, and increasingly in Africa, where production has been steadily growing in recent years (Haverkort and Struik, 2015). However, despite its widespread cultivation, potato production is highly vulnerable to various pests and diseases, which significantly affect yield and quality. The most devastating threaten is late blight, caused by the oomycete pathogen *Phytophthora infestans* (Mont.) de Bary. Late blight can affect all parts of the plant, including stems, leaves, and tubers, leading to severe crop losses and threatening food security (Fry, 2008). The economic costs associated with late blight, including crop loss and chemical control, exceed €9 billion annually (Haverkort et al., 2016). Historically, *P. infestans* caused widespread famine in Europe, particularly in Ireland, underscoring the catastrophic potential of this disease (Zadoks, 2008). Given the global importance of potatoes as a food source, addressing the challenges posed by late blight is critical. One promising avenue for sustainable disease management is the utilization of disease resistance genes in potato breeding programs. This approach offers a long-term solution to mitigate the impact of late blight, reduce chemical use, and enhance crop productivity.

To date, over 70 *R* genes, encoding nucleotide-binding domain (NB) leucine-rich repeat (LRR)-containing receptors (NLRs), have been identified or cloned from wild potato relatives (Paluchowska et al., 2022). However, the emergence and rapid evolution of new virulent strains of *P. infestans* have led to the breakdown of resistance in many of these *R* genes (Paluchowska et al., 2022; Vleeshouwers et al., 2011). Despite this, *Rpi-vnt1.1*, derived from *S. venturii* (Foster et al., 2009), has shown strong resistance against field isolates of *P. infestans* in China, outperforming other *R* genes, including *RB* and *R8* (Zhang et al., 2025). However its potential, this gene has not been widely incorporated into commercial potato cultivars (Paluchowska et al., 2022). Therefore, developing and deploying this resistance gene in the field is crucial for controlling late blight and improving potato yield.

To detect the presence of *Rpi-vnt1.1* in advanced breeding lines from the International Potato Center (CIP), we exploited pan-genomic data (Tang et al., 2022) to analysis gene-specific sequences. We collected 440 homologous gene sequences of *Rpi-vnt1.1* from the potato pan-genome, including known homologs *Rpi-vnt1.2, Rpi-vnt1.3, Tm22*, and *ph3* (Paluchowska et al., 2022). Simulating the design principles of probe primers, we fragmented the sequences of these 440 genes into 20 bp short-reads and remapped them to the sequence of *Rpi-vnt1.1* (Table S1). Our analysis revealed that non-functional *Rpi-vnt1.1* homologs lack a complete 5′-end extension that coding the first 33 amino acids, whereas functional alleles conferring resistance carry the intact 5′-end extension (Figure 1A). This observation enabled us to design a specific molecular marker targeting the 5′-end extension region for screening potato germplasm harboring functional *Rpi-vnt1.1* allele. The primer sequences are shown in Figure 1A and Table S2.

**Figure 1.**
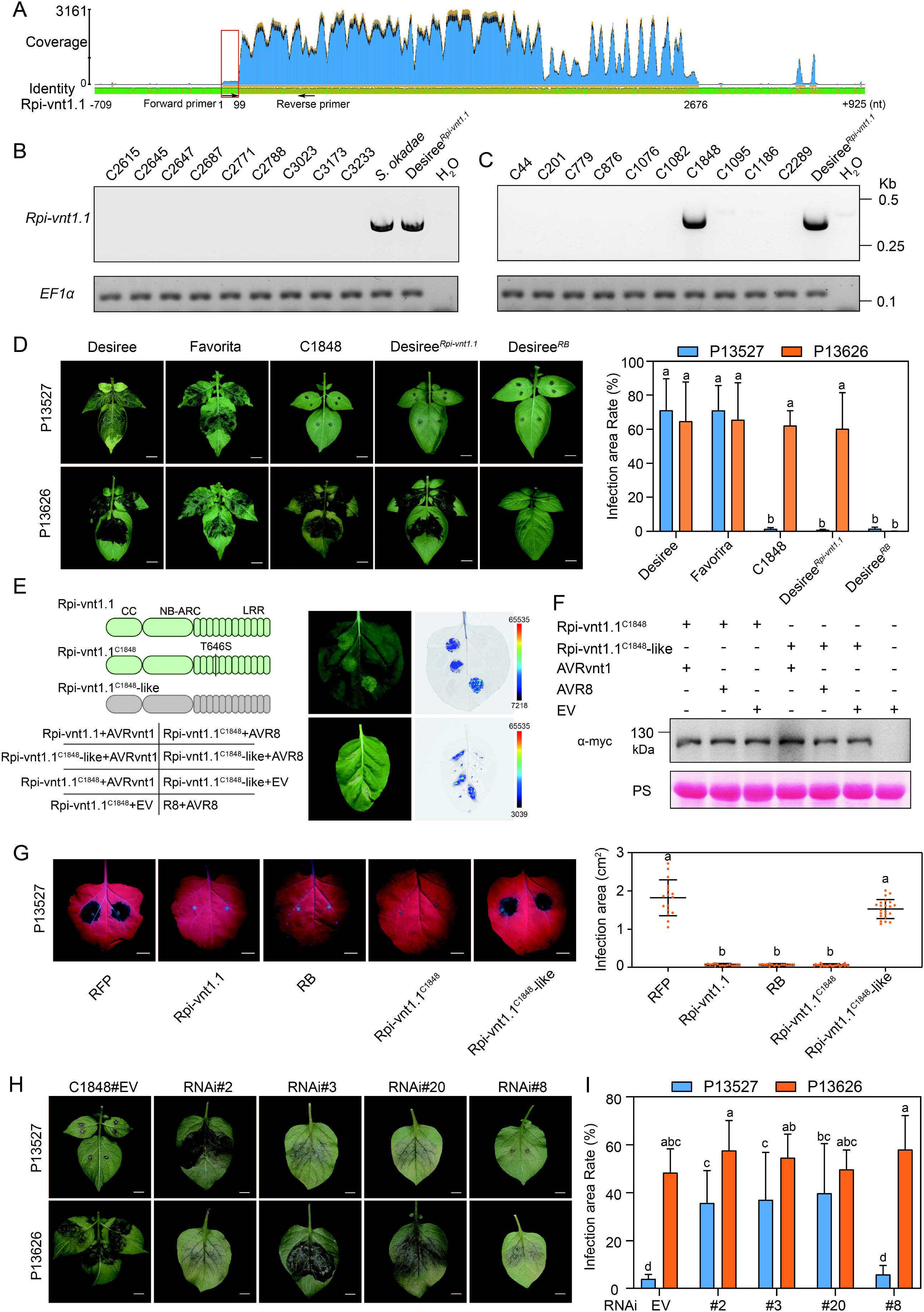
Design molecular marker for identifying functional *Rpi-vnt1.1* homologs in potato resources. (**A**) Design the molecular marker for *Rpi-vnt1.1* based on potato Pan-genome. The red box indicates the specific 5′-end extension sequence unique to functional *Rpi-vnt1.1* alleles. The arrows denote the primer locations. (**B**) and (**C**) Specificity testing of the functional *Rpi-vnt1.1* marker was performed by PCR amplification using potato genomic DNA from wild species and advanced breeding lines, respectively. The potato *EF1α* gene was used to as a positive control. Desiree^*Rpi-vnt1.1*^ represents the transgenic potato Desiree carrying *Rpi-vnt1.1*, and”H_2_O”is the negative control. (**D**) Evaluation of disease resistance in advanced breeding line C1848. Cultivars Desiree and Favorita were used as susceptible negative controls. Desiree^*Rpi- vnt1.1*^ and Desiree^*RB*^ represent transgenic potato lines generated in the Desiree background. Leaves from 4-week-old potato plants were inoculated with *P. infestans* isolates P13527 and P13626. P13626, an *AVRvnt1*-silenced isolate, overcomes *Rpi-vnt1.1*-mediated resistance, whereas P13527 cannot. Infected potato leaves were photographed at 5 d post-inoculation (dpi) (left) and the infection area rate was estimated by measuring the lesion area (right). The data are shown as means ± SD (n = 3). “a” and “b” denote statistically significant differences according to one-way analysis of variance followed by Tukey’s test (*P* < 0.05). Scale bar, 10 mm. (**E**) Diagram showing sequences of *Rpi-vnt1.1, Rpi-vnt1.1*^*C1848*^ and *Rpi-vnt1.1*^*C1848*^*-like*. Co-expression of the *Rpi-vnt1.1*/*Rpi-vnt1.1*^*C1848*^ construct with *AVRvnt1* induces HR in *N. benthamiana* leaves at 5 dpi. HR induced by *Rpi-vnt1.1*/*AVRvnt1* and *R8*/*AVR8* combinations served as positive controls. *Rpi-vnt1.1*^*C1848*^*-like*/*AVRvnt1, Rpi-vnt1.1*^*C1848*^*-like*/*AVR8, Rpi-vnt1.1*^*C1848*^/*AVR8, Rpi-vnt1.1*^*C1848*^/EV, and *Rpi-vnt1.1*^*C1848*^*-like*/EV combinations served as negative controls. EV, empty vector. (**F**) Immunoblot analysis was performed using anti-myc antibodies to detect *Rpi-vnt1.1*^*C1848*^-myc and *Rpi-vnt1.1*^*C1848*^*-like*-myc proteins under various combination conditions, as shown in panel (**E**). Total protein was extracted from *N. benthamian*a leaves at 24 h post-inoculation (hpi). Protein loading is shown by Ponceau S staining for Rubisco (RBC). (**G**) Expression of *Rpi-vnt1.1*^*C1848*^, *Rpi-vnt1.1* and *RB* genes, but not *RFP* and *Rpi-vnt1.1*^*C1848*^*-like*, confers resistance to *P. infestans* isolate P13527 infection. *N. benthamiana* leaves expressing *RFP* or *R* genes were inoculated with *P. infestans* zoospore suspension at 24 hours after *Agrobacterium* infiltration. Infected leaves were photographed under UV light (left panel) and lesion areas were measured (right panel). The data are shown as means ± *SD*. ‘a’ and ‘b’ denote statistically significant differences according to one-way ANOVA followed by Tukey’s test (*P* < 0.05). (**H**) Zoospore suspension of *P. infestans* strain P13527 or P13626 was drop-inoculated on detached leaves of C1848#EV and C1848 RNAi transgenic potato lines. The infected leaves were photographed at 5 dpi. (**I**) Infected areas rate (%) of inoculated leaves in (**H**). The infected area was measured in ImageJ. The data represent the mean±SD from indicated replicates. Different letters denote a significant difference at *P* < 0.05 using one-way ANOVA with Tukey’s HSD test.

To evaluate the specificity of the designed marker, we tested ten accessions each from wild relatives, landraces, and breeding lines. The marker did not amplify the Rpi-vnt1.1-specific fragment in any of these accessions. As expected, the fragment was successfully amplified in both the wild diploid *S. okadae*, which carries a functional *Rpi-vnt1.1* gene (Van Weymers et al., 2016), and the positive control (genomic DNA from transgenic Desiree carrying *Rpi-vnt1.1*) (Figures 1B, S1). Notably, amplification was successful in one advanced breeding line, C1848 (CIP392032.2), confirming the presence of *Rpi-vnt1.1* (Figure 1C). These results demonstrate that the 5′-end extension-based primers are highly specific and enabled the identification of a potato line carrying the broad-spectrum resistance gene *Rpi-vnt1.1*.

To assess the resistance level of C1848 against late blight, we inoculated C1848, two susceptible cultivars (Desiree and Favorita), and transgenic Desiree lines carrying either *Rpi-vnt1.1* or *RB*, using the *P. infestans* isolate P13527 (Pais et al., 2018). C1848 and the transgenic *Rpi-vnt1.1* and *RB* lines exhibited strong resistance, while Desiree and Favorita were susceptible. When inoculated with the *AVRvnt1*-silenced strain P13626 (Gao et al., 2020; Pais et al., 2018), C1848 and the *Rpi-vnt1.1* transgenic line, as well as Desiree and Favorita, were susceptible, whereas *RB* transgenic line retained resistance (Figure 1D). These results indicate that late blight resistance of C1848 is specifically mediated by functional *Rpi-vnt1.1*.

To further confirm the sequence of *Rpi-vnt1.1* in C1848, we cloned the gene using specific primers (Table S2). Sequence analysis identified two variants: one allele with a single amino acid substitution (T646S) compared to the canonical *Rpi-vnt1.1*, designated *Rpi-vnt1.1*^*C1848*^, and another with only 75% sequence identity, designated *Rpi-vnt1.1*^*C1848*^*-like* (Figure 1E). To determine whether the T646S substitution affects gene function, we transiently co-expressed each allele with cognate effector AVRvnt1.1 in *Nicotiana benthamiana*. Only *Rpi-vnt1.1*^*C1848*^ triggered a hypersensitive response (Figure 1E). Furthermore, overexpression assays in *N. benthamiana* leaves revealed that *Rpi-vnt1.1*^*C1848*^, the canonical *Rpi-vnt1.1*, and *RB* conferred resistance against isolate P13527, whereas *Rpi-vnt1.1*^*C1848*^*-like* did not (Figure 1G). Protein accumulation assays confirmed that both *Rpi-vnt1.1*^*C1848*^ and *Rpi-vnt1.1*^*C1848*^*-like* were stably expressed (Figure 1F). These findings demonstrate that the T646S substitution does not compromise the resistance function of *Rpi-vnt1.1*.

To provide genetic evidence that *Rpi-vnt1.1*^*C1848*^ alone confers resistance in C1848, we employed RNAi-mediated gene silencing. Three independent transgenic RNAi lines (RNAi#2, RNAi#3, RNAi#20) displayed compromised resistance when inoculated with P13527, whereas RNAi#8 and the empty vector control (C1848#EV) remained resistant (Figures 1H, S2). The qRT-PCR analysis showed markedly reduction of *Rpi-vnt1.1*^*C1848*^ transcript levels in RNAi#2, RNAi#3, and RNAi#20 compared to RNAi#8 and C1848#EV, correlating with the observed loss of resistance (Figure 1I, S3). Collectively, these results provide genetic confirmation that resistance in C1848 is majorly mediated by the functional allele *Rpi-vnt1.1*^*C1848*^.

During sequence analysis and resistance phenotyping of *Rpi-vnt1.1*, we made the following key observations: (1) interrogation of a potato pan-genome NLR database revealed a distinctive 5′-end extension that is present in functional *Rpi-vnt1.1* alleles but absent from non-functional homologues; (2) exploiting this N-terminal signature, we designed a simple, low-cost and rapid molecular marker for *Rpi-vnt1.1*, and validated its specificity across a panel of germplasm; and (3) using this marker to screen advanced breeding lines from the CIP, we identified a breeding line, C1848, that carries a functional *Rpi-vnt1.1* allele. Functional assays confirmed that the *Rpi-vnt1.1* allele in C1848 confers broad-spectrum resistance to *P. infestans*. Given the limited deployment of *Rpi-vnt1.1* in widely grown cultivars, the 5′-end extension–based marker provides a practical tool for marker-assisted selection and rapid identification of resistant material in breeding programs. Together, these findings facilitate the targeted deployment of *Rpi-vnt1.1* to enhance the durability of late blight resistance and improve potato yield stability.

*Rpi-vnt1.1* was cloned from the diploid wild species *S. venturii* and confers a broad-spectrum resistance against *P. infestans* (Foster et al., 2009). Zhang et al. showed that *Rpi-vnt1.1* consistently confers strong resistance (Zhang et al., 2025), implying that its cognate AVR effector, AVRvnt1.1, is conserved across most *P. infestans* populations — with the notable exception of isolate P13626 (Pais et al., 2018). In our study, inoculation with an *AVRvnt1*-silenced *P. infestans* strain P13626 abolished resistance in potato C1848, demonstrating that the observed resistance in C1848 is specifically mediated by functional *Rpi-vnt1.1* allele.

Cultivated potato is a highly heterozygous, auto-tetraploid crop, which complicates reliable PCR-based detection of known resistance genes because of allelic variation, paralogues and copy-number complexity (Wu et al., 2023). Diagnostic resistance gene enrichment Sequencing (dRenSeq) can accurately identify known *R* genes in such complex genomes (Armstrong et al., 2019), but it is substantially more expensive than PCR. In this study, by mining NLR sequence features we discovered a distinctive N-terminal signature in functional *Rpi-vnt1.1* alleles, which was used for developing a PCR-based molecular marker. This marker is low-cost, rapid, and specific (as validated across germplasm), making it a practical and scalable tool for identifying *Rpi-vnt1.1* containing germplasms and for marker-assisted selection in late blight breeding programs.

Although *Rpi-vnt1.1* was first identified in 2009, it has not been widely deployed in tetraploid cultivated potato—largely because introgression of resistance genes from diploid wild relatives into tetraploid cultivars is frequently constrained by crossing barriers, differing endosperm balance numbers and protracted breeding cycles. In this study, we applied a specific 5′-end extension–based molecular marker to screen tetraploid advanced lines and identified the advanced breeding line C1848, which carries a functional *Rpi-vnt1.1* allele. The availability of this tetraploid material provides an important genetic resource that will facilitate incorporation of *Rpi-vnt1.1* into breeding programs and enable its broader deployment for durable late blight resistance in cultivated potato.

## Materials and methods

### Plant materials and *P. infestans* infection assays

A total of 40 potato accessions were used in the study, including cultivars, breeding lines, wild species, and landraces (Table S3). Potato and *N. benthamiana* plants were grown and maintained in a greenhouse at 22–25°C, 50% relative humidity, and 75–100 mE m^−2^ s^−1^ light with a 12-h light/12-h dark photoperiod (Zhao et al., 2025). *P. infestans* isolates were grown on rye sucrose agar (RSA) media plates for 9–12 d at 18°C as described elsewhere (Song et al., 2009). Sporangia were harvested from plates using cold water and zoospores were collected 1–2 h after incubation at 4°C (Huang et al., 2020). Zoospore suspension of *P. infestans* isolates were inoculated on 3–4 weeks old detached potato or *N. benthamiana* leaves, and disease symptoms were assessed at 4–5 dpi.

### Analysis of *Rpi-vnt1.1* sequence

Homologs of *Rpi-vnt1.1* from the potato pan-genome were previously identified using the ‘NLR local annotation’ pipeline based on 44 high-quality diploid potato genomes (Tang et al., 2022). A total of 440 *Rpi-vnt1.1* homolog sequences were extracted and fragmented into 20 bp short reads using the ‘sliding’ function in seqkit v2.10 with parameters ‘-s 10 -w 20’ (Table S3). These short reads were then converted to FASTQ format using the FASTA_to_FASTQ.py script from the FASTAX-Toolkit v0.0.13. All short reads were mapped to the reference *Rpi-vnt1.1* full-length sequence using minimap2 v2.2.30 with the parameter ‘-ax sr’. The resulting SAM file was sorted into a BAM file with samtools v1.22.1 and imported into Geneious Prime v2025.2 for visualization of the mapped reads.

### Transformation and RNAi of potato

For hairpin RNA interference (RNAi) targeting *Rpi-vnt1.1*^*C1848*^, a fragment containing the chalcone synthase (CHSA) intron was amplified from the plasmid *pFGC5941*. PCR-derived 5′-*Rpi-vnt1.1*^*C1848*^ cDNA fragments were designed to overlap with the 5′ and 3′ ends of the CHSA intron in opposite orientations (Tables S2, S4). The resulting fusion fragment was cloned into the *pCAMBIA1300* vector, which was then introduced into *Agrobacterium tumefaciens* strain GV3101 for subsequent potato transformation (Sun et al., 2024). For generating potato transgenic plants, 3-week-old potato plantlets C1848 gown in MS medium (Zhao et al., 2025) in a growth incubator at 20–23°C, 50% relative humidity, and 75–100 mE m-2 s^−1^ light with a 12-h light/12-h dark photoperiod were used for *A. tumefaciens*-mediated plant transformation as previously described with minor modifications (Ducreux et al., 2005). After a 2-d pre-culture period, the explants were co-cultured with *Agrobacterium* strain GV3101 carrying the RNAi constructs for another 2 d. During this co-cultivation, 2 mg/L of α-naphthaleneacetic acid and 1 mg/L of trans-zeatin were added. Subsequently, α-naphthaleneacetic acid (0.01 mg/L) and trans-zeatin (2 mg/L) were used to promote callus formation and re-generation until visible shoots appeared, and the positive transformants were selected on kanamycin (50 mg/L). Gene silencing transgenic plants were further confirmed by RT-qPCR.

### Total RNA isolation and RT-qPCR analysis

Total RNA was extracted from leaves of 6-week-old transgenic potato plants using the Super FastPure Cell RNA Isolation Kit (RC102, Vazyme Biotech Co.,Ltd, China) and quantified with a NanoDrop spectrophotometer (Thermo Fisher Scientific, USA). One microgram of RNA was treated with gDNA wiper Mix (Vazyme Biotech Co.,Ltd, China) at 42 °C for 2 minutes to remove genomic DNA, followed by reverse transcription using HiScript II qRT SuperMix II (Vazyme Biotech Co.,Ltd, China) at 50 °C for 15 minutes. Quantitative real-time PCR was carried out on a 7500 Fast Real-Time PCR System (Applied Biosystems, USA) using the HiScript II One Step qRT-PCR SYBR Green Kit (Vazyme Biotech Co.,Ltd, China) with gene-specific primers listed in Table S2. Gene expression levels were normalized to the internal reference gene *EF1α*.

### Cloning of *Rpi-vnt1.1*^*C1848*^ and Agroinfiltrations

The homologous sequence of *Rpi-vnt1.1* was amplified from potato C1848 genome and cloned into pBin308 vector using ClonExpress Ultra One Step Cloning Kit V3 (C117, Vazyme Biotech Co.,Ltd, China). Construct plasmids were transformed into *A. tumefaciens* strain GV3101 and grown overnight in Luria-Bertani (LB) culture medium with the antibiotic kanamycin and rifampicin at 28°C. Transformed bacteria were harvested by centrifugation at 5000×g for 4 min and then resuspended and diluted in agroinfiltration buffer (10 mM MgCl_2_, 150 mM acetosyringone, 10 mM MES) to an optical density at 600 nm (OD_600_) ranging from 0.01 to 1.0, depending on experimental requirements. After incubation in the dark at room temperature for 2 to 4 hours, the bacterial suspensions were infiltrated into the abaxial side of leaves from 4-to 5-week-old *N. benthamiana* plants using a needleless syringe. For co-expression of multiple constructs, suspensions containing each construct were mixed thoroughly prior to infiltration.

## Author contributions

Xiaoqiang Zhao conceived the research. Xiaoqiang Zhao, Shuo Wang, Simeng Feng, and Chenxi Hu performed the experiments. Luyao Wang analyze the data. Shiwei Chang and Yan Feng provided and assessed the experimental materials. Shuo Wang and Xi Meng performed the phenotypic assessment. Xiaoqiang Zhao wrote the manuscript. Liang Kong revised the manuscript. All authors read and approved the final manuscript.

## Acknowledgements

We thank Dr. Jack H. Vossen (Wageningen University) for providing the transgenic potato lines carrying *RB* and *Rpi-vnt1.1*. We thank the Bioinformatics Center of Nanjing Agricultural University for their support. This research was financially supported by Hebei key Research and Development Program (Grant No. 21326320D), Science and Technology Plan Projects in Inner Mongolia Autonomous Region (No. 2024KJHZ0005) and the National Natural Science Foundation of China (No. 32130088).

## Figure legends

**Figure S1.**
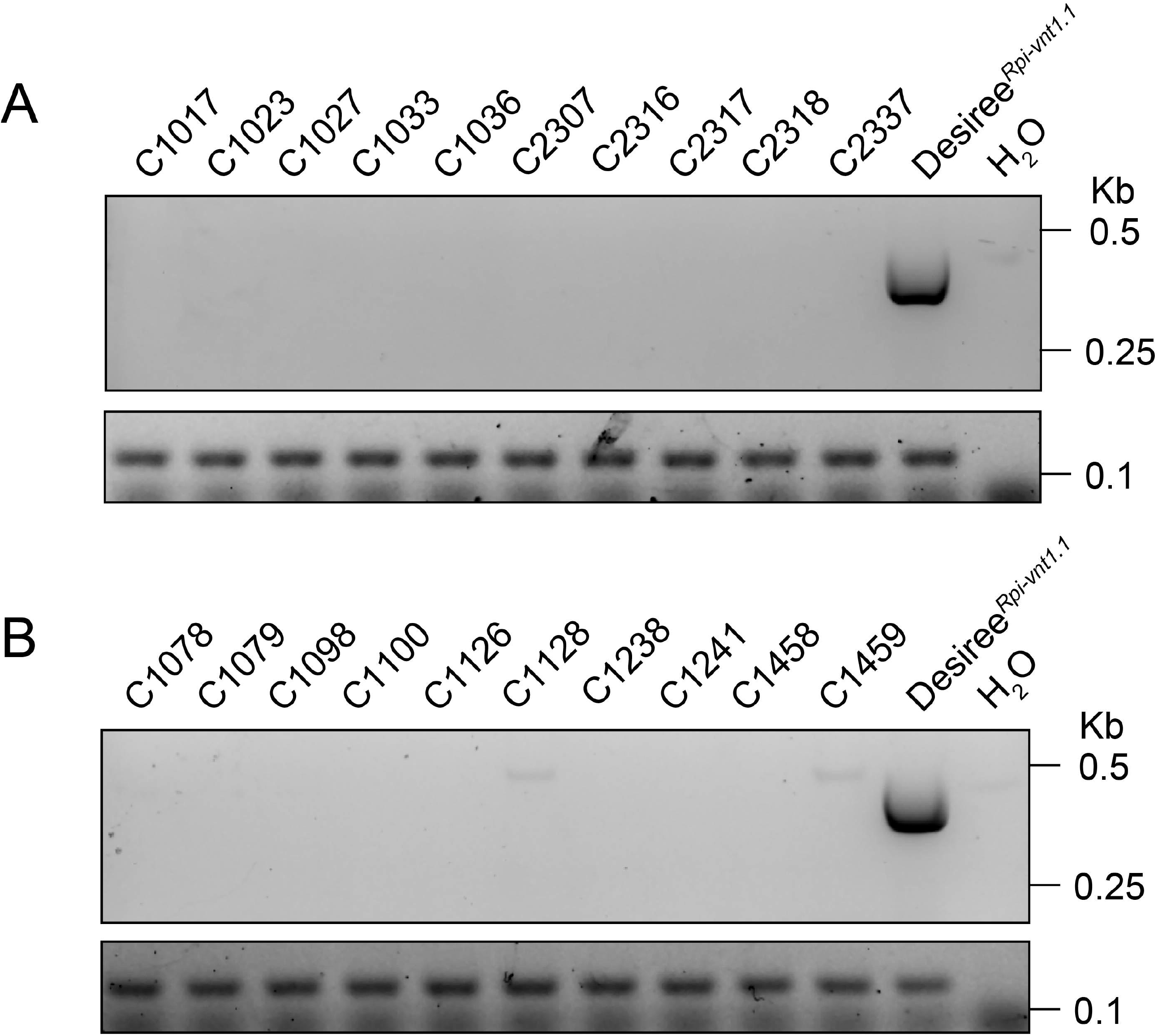
Specificity testing of the functional *Rpi-vnt1.1* marker was performed by PCR amplification using potato genomic DNA from native landraces (**A**) and breeding lines (**B**), respectively.

**Figure S2.**
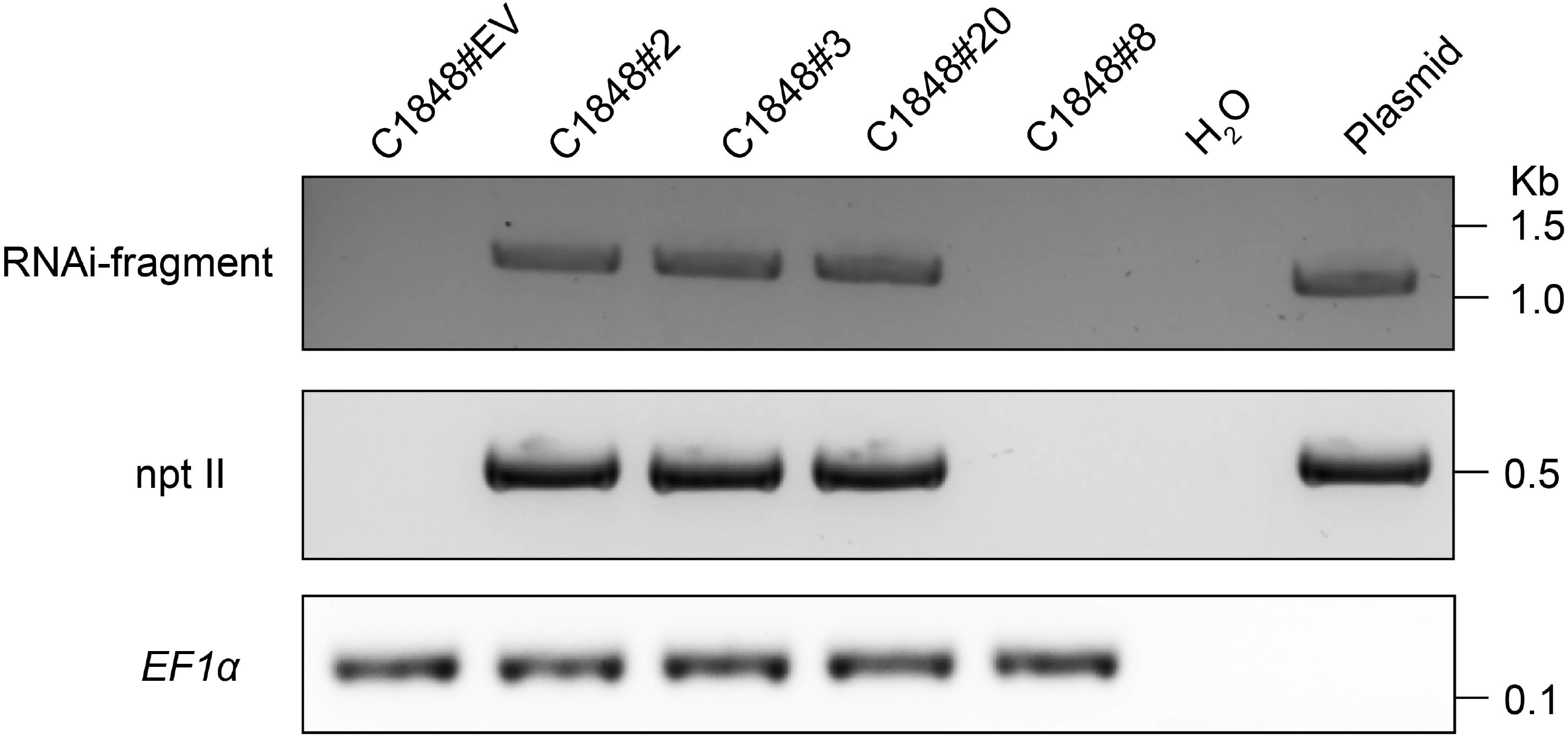
Identification of C1848 RNAi transgenic Potato Plants by polymerase chain reaction (PCR)-genotyping. Genotyping PCR, using genomic DNA, was performed with primers listed in Table S2 for screening C1848 RNAi transgenic Potato. The potato *EF1α* gene was used to as a positive control. “Plasmid” denotes the plasmid DNA control, while “C1848#EV” represents the transgenic line transformed with the empty vector, which carries no silencing fragment. “npt II” serves as the selectable marker for screening T-DNA insertion, and “H_2_O” is included as the negative control.

**Figure S3.**
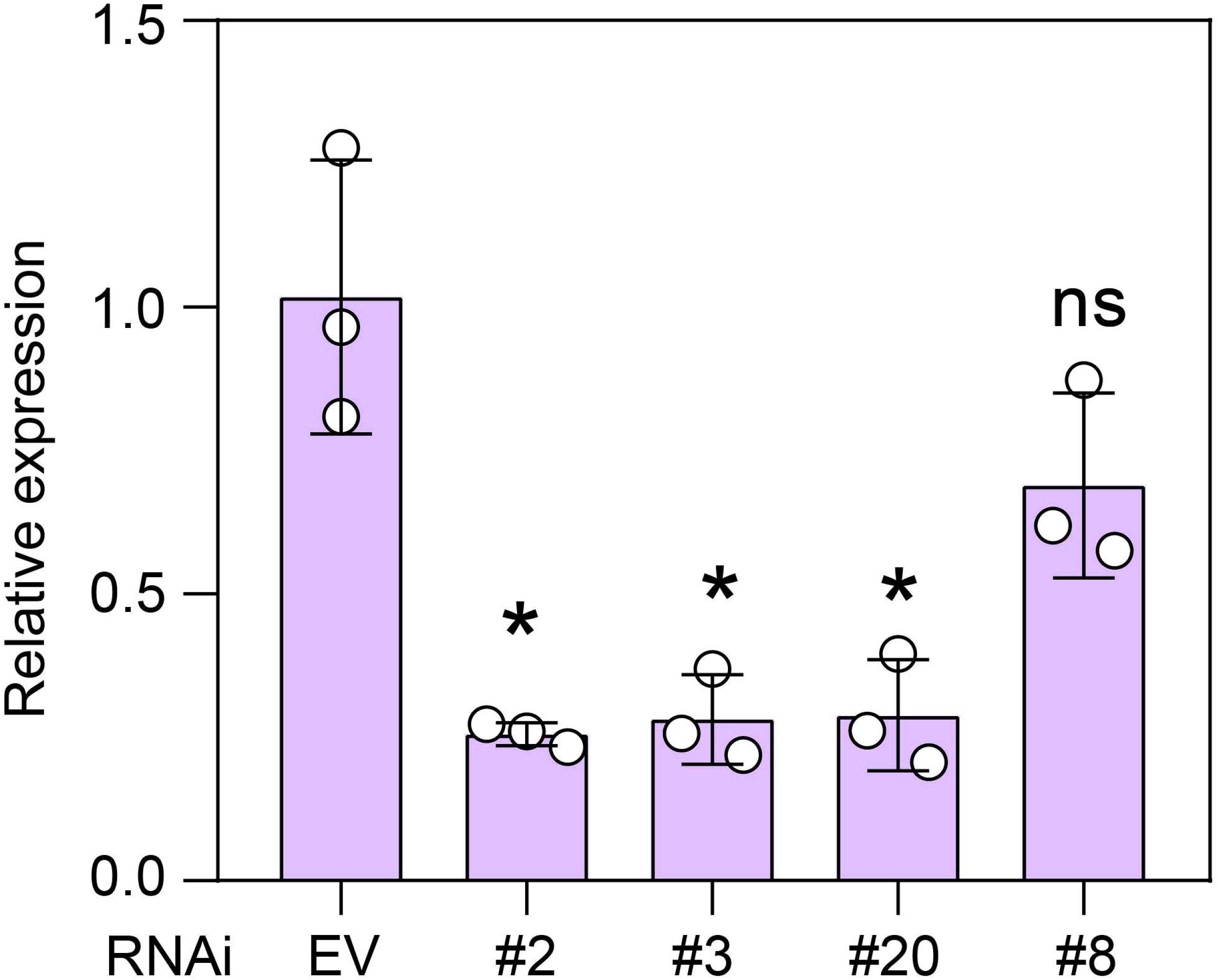
Relative expression of *Rpi-vnt1.1*^*C1848*^ in 3 RNAi transformed lines compared to C1848#EV transgenic potato. Bars represent the mean ± SD of 3 independent replicates. Asterisks designate a statistical difference from the C1848#EV control at *P* < 0.05 using one-way ANOVA with Tukey’s test.

